# A fiducial-based framework for precise MRI-guided stereotaxic targeting in nonhuman primates

**DOI:** 10.64898/2026.08.28.747848

**Authors:** Peyton Harmon, Reza Azadi

## Abstract

**Background:** Accurate MRI-guided stereotaxic targeting in nonhuman primates requires reliable registration between MRI coordinates and stereotaxic space. We evaluated skull-fixed MR-compatible screw fiducials as a practical reference for this registration.

**Results:** Using a 3D-printed skull model with 28 fiducials and three internal targets, we compared offset translation, rigid registration, and affine registration under aligned and misaligned conditions. Offset translation was most sensitive to misalignment. Affine registration fit skull-surface fiducials closely, but rigid registration produced lower overall target errors. In vivo validation showed that implanted fiducials could be identified on MRI and re-measured during a later surgery.

**Comparison with existing methods:** This approach builds on previous fiducial-based stereotaxic methods by systematically comparing registration transforms, testing misalignment, evaluating fiducial number, and measuring accuracy at independent internal targets.

**Conclusions:** Skull-fixed fiducials provide a practical framework for MRI-to-stereotaxic registration. Rigid registration may offer the safest balance between correcting positioning differences and preserving accuracy at internal targets.

**Highlights:**

- MR-compatible screws provide an MRI-to-stereotaxic registration framework
- Offset translation is sensitive to stereotaxic misalignment
- Rigid registration improves accuracy at independent internal targets
- Affine registration can overfit skull-surface fiducials
- Implanted screw fiducials supported in vivo stereotaxic registration

## 1. Introduction

Nonhuman primates (NHPs) are important models for neuroscience research because of the anatomical and functional similarities between their brains and those of humans. Many experimental approaches in NHPs, including electrophysiological recordings, microinjections, viral vector delivery, and lesion studies, require precise localization of cortical and subcortical structures. Stereotaxic techniques have long provided a framework for targeting these structures using a three-dimensional coordinate system referenced to anatomical landmarks Horsley and Clarke (1908). Accurate targeting is particularly important for small or deep brain structures, where relatively small localization errors can affect experimental outcomes.

For MRI-guided stereotaxic procedures, coordinates measured in image space must be related to coordinates in the stereotaxic frame. Historically, this correspondence has been established using MRI-visible earbars Saunders, Aigner and Frank (1990). Other approaches have used anatomical landmarks, such as the sagittal sinus McBride and Clark (2016), or external fiducial markers Liang, Zimmermann Rollin, Alikaya, Ho, Santini, Bostan, Schwerdt, Stauffer, Ibrahim, Pirondini and Schaeffer (2024). Registration based on a limited number of landmarks, however, can remain sensitive to differences in positioning between imaging and stereotaxic procedures, potentially reducing targeting accuracy.

Any feature that can be identified both in preoperative imaging and during stereotaxic procedures can serve as a fiducial reference for registration between image and physical coordinate systems. Multiple fiducials distributed across the skull may provide a more robust basis for registration than a small number of anatomical landmarks. Fiducial-based registration is widely used in clinical neurosurgery and image-guided procedures, and related approaches have been described in nonhuman primates using skull-mounted MRI-visible fiducials for stereotaxic targeting Bentley, Khalsa, Kobylarek, Schroeder, Chen, Bergin, Tat, Chestek and Patil (2018). However, the effects of registration model, stereotaxic misalignment, and fiducial number on MRI-to-stereotaxic registration accuracy remain incompletely characterized.

In this study, we developed and evaluated a skull-fixed fiducial-based framework for registering MRI coordinates to stereotaxic space in nonhuman primates. Using fiducials that could be identified both on MRI and during stereotaxic measurement, we established direct correspondence between imaging and physical coordinate systems and quantified registration accuracy using independent internal targets. We compared three registration approaches with increasing degrees of flexibility: offset translation, which applies a uniform coordinate shift; rigid registration, which allows rotation and translation while preserving distances; and affine registration, which additionally allows scaling and shear. We further evaluated the robustness of each approach following stereotaxic misalignment and examined the effect of fiducial number on registration performance. These analyses provide a practical framework for evaluating and improving MRI-guided stereotaxic targeting in nonhuman primates.

## 2. Methods

### 2.1. 3D-printed model and MRI scan

#### 2.1.1. Materials

- MRI-compatible stereotactic head frame for nonhuman primates; Jerry-Rig
- Micromanipulator; Kopf
- 3D-printed skull model
- Agar powder
- Gadolinium-diethylenetriamine pentaacetic acid (Gd-DTPA); Magnevist

#### 2.1.2. Methods

A 3D-printed skull model was used to evaluate the accuracy of MRI-to-stereotaxic registration (Figure 1A;Supplementary File 1). Twenty-eight MR-compatible screw fiducials were placed on the skull surface. Three additional markers were placed along the midline inside the skull model to serve as internal targets that were not used to estimate the registration transformations. To provide MRI-visible contrast around the fiducials and internal targets, the model was embedded in an 8% agar solution containing 5 mmol/L Gd-DTPA and allowed to solidify (Figure 1B,C).

**Figure 1.**
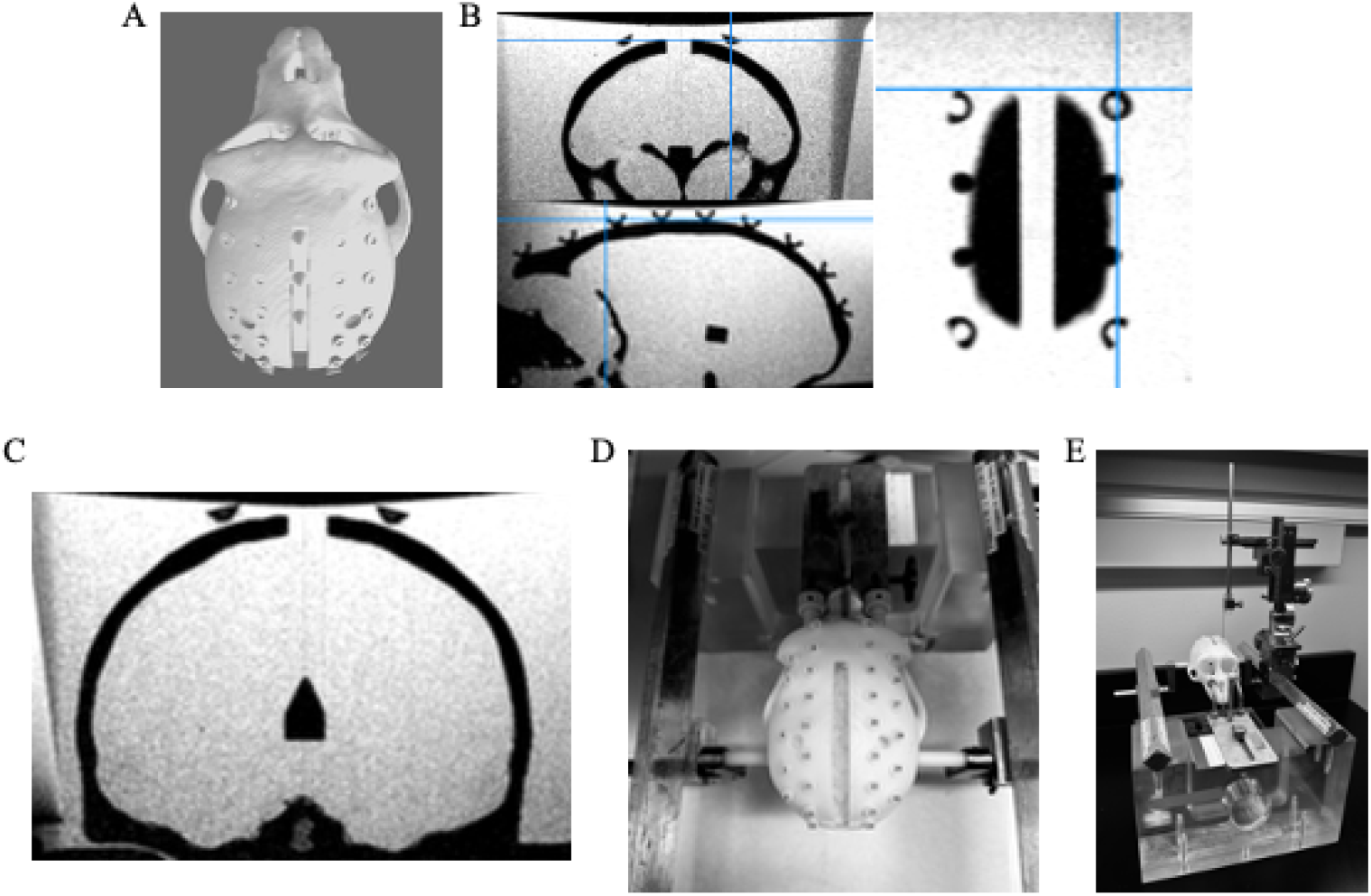
Skull model and fiducial measurement procedure. (A) Skull model with screws placed on the skull surface to serve as fiducial markers. (B) MRI scan of the 3D-printed skull model embedded in agar containing contrast agent, showing the skull-surface screw fiducials. (C) MRI section showing the internal midline targets. (D) 3D-printed skull model mounted in the stereotaxic frame for direct coordinate measurement. (E) Fiducial coordinates were measured in stereotaxic space using a stereotaxic micromanipulator.

MRI was performed with the model mounted in an MRI-compatible stereotaxic frame. The coordinates of each fiducial and internal target were measured in the MRI volume and then measured directly on the physical model using a stereotaxic micromanipulator. For the skull-surface fiducials, the most anterior point of each screw head was used as the fiducial reference point in both coordinate systems. Each stereotaxic measurement was repeated three times in random order, and the average coordinate was used for subsequent analyses (Figure 1D,E).

To evaluate the robustness of the registration methods to framing errors, stereotaxic measurements were repeated after intentionally misaligning the model within the stereotaxic frame. This was done by removing the right earbar from the ear canal before re-measuring the fiducial and target positions. This condition is referred to as the misaligned condition in the Results.

### 2.2. In Vivo Surgical Validation

#### 2.2.1. Materials

- MRI-compatible stereotaxic head frame for nonhuman primates; Jerry-Rig
- Ceramic bone screws, 5 mm; Thomas Recording
- Carbide burs, HP-7; SS White Dental
- High-speed surgical drill set; BASi MF-5360
- Screwdriver; Thomas Recording
- Taps for ceramic screws; Thomas Recording
- Stereotaxic micromanipulator; Kopf

#### 2.2.2. Methods

The in vivo feasibility of the fiducial-based registration approach was evaluated in a rhesus monkey across two surgical procedures. All surgical procedures were performed under sterile conditions and general anesthesia, following the surgical procedures described in Tangen, Harmon, Plotnikova, Mohanty, Cummins, Pelkey, Cameron, Rood, Guerriero, McBain, Eldridge, Averbeck and Azadi (2026). One twelve-year-old male monkey (*Macaca mulatta*) was used in this study. All experimental procedures adhered to the Guide for the Care and Use of Laboratory Animals and were reviewed and approved by the National Institute of Mental Health (NIMH) Animal Care and Use Committee.

During the first surgery, the skin and fascia were opened and the muscle was reflected to expose the skull. Ten ceramic MR-compatible screws were implanted as cranial fiducials, with five screws placed over each hemisphere. The screw locations were spaced approximately 5 mm apart. For each fiducial, a hole was drilled using a high-speed surgical drill with irrigation to reduce heat-related bone damage. Each hole was then tapped, and the screw was inserted manually using the matched screwdriver with slow, controlled rotation. After screw implantation, the muscle, fascia, and skin were closed over the fiducials, and the animal was allowed approximately three weeks to recover.

After recovery and before the second surgery, the animal was positioned in the MRI-compatible stereotaxic frame and scanned. Three T1-weighted MRI scans were acquired and averaged to improve localization accuracy. The implanted screws were identified in the MRI volume, and the most anterior point of each screw head was recorded as the fiducial reference point, as in the 3D-printed model (Figure 3A,B). These MRI fiducial coordinates were then used for registration to stereotaxic space.

**Figure 2.**
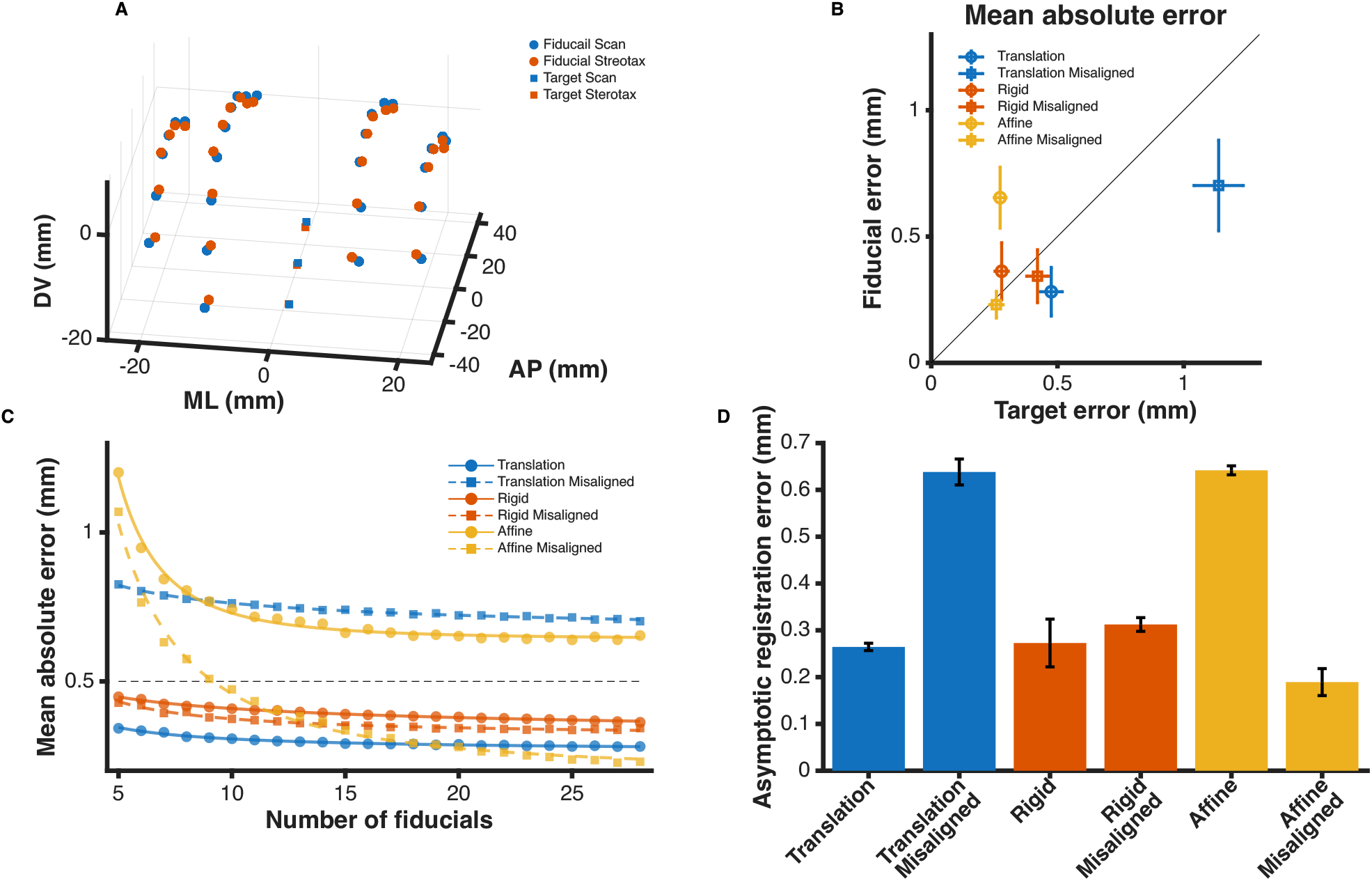
Fiducial-based registration accuracy in the 3D-printed skull model. (A) Three-dimensional distribution of skull-surface fiducials and internal targets measured in MRI scan space and stereotaxic space. Fiducials were distributed across the skull surface (circles), whereas the internal targets were positioned along the midline and were not used to estimate the registration transformations (squares). (B) Relationship between target error and fiducial error for offset translation (blue), rigid registration (red), and affine registration (yellow) under aligned (circles) and misaligned (squares) conditions. The diagonal line indicates equal fiducial and target errors. (C) Mean absolute coordinate error as a function of the number of fiducials used to estimate each transformation. Errors decreased with increasing fiducial number and approached an asymptotic minimum. The black dashed line indicates the 0.5-mm MRI voxel size. (D) Estimated asymptotic target registration error for each method and alignment condition, obtained from power-law fits to the fiducial-number curves. Error bars indicate 95% confidence intervals.

**Figure 3.**
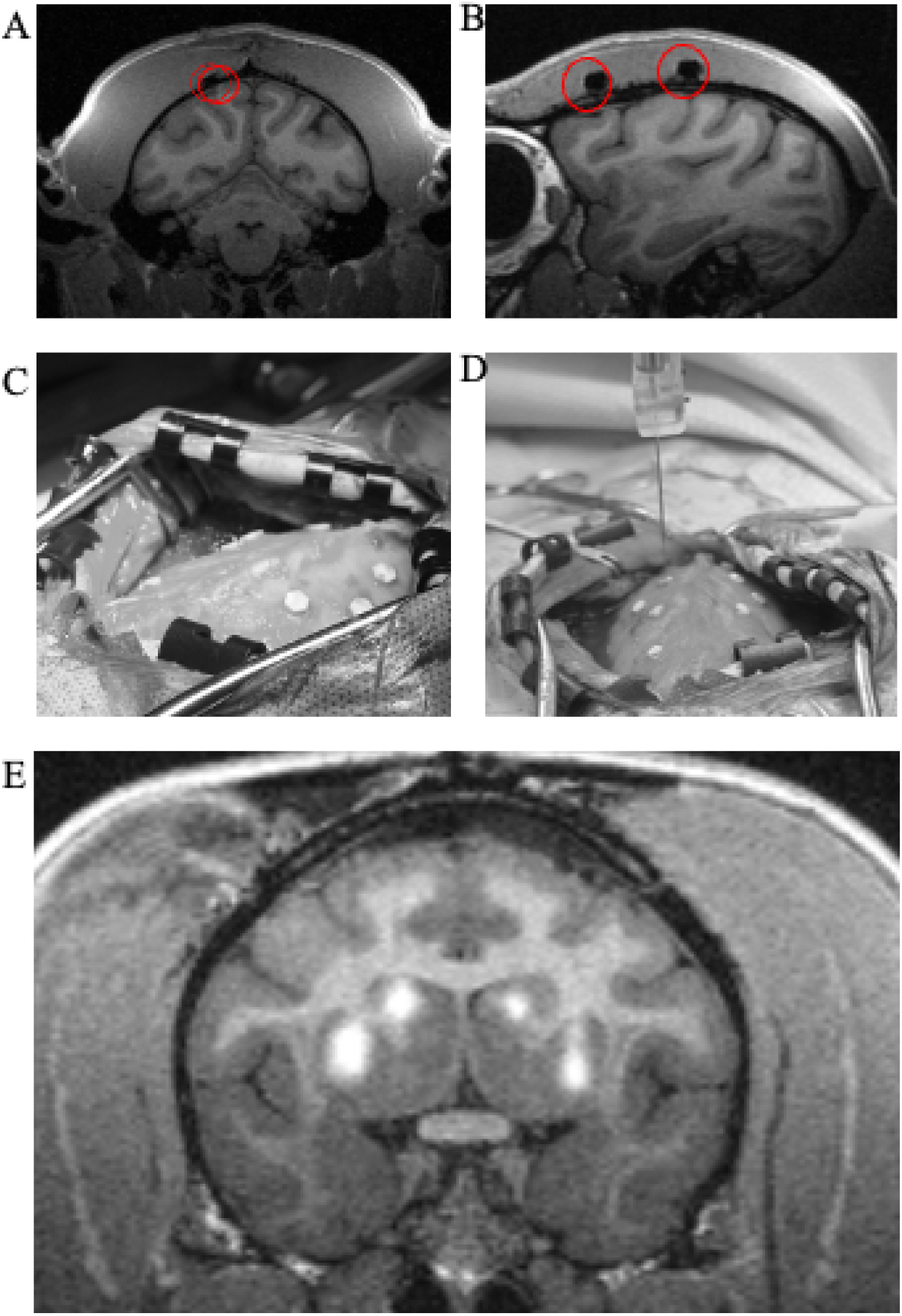
In vivo fiducial implantation and stereotaxic measurement. (A, B) MRI sections obtained after implantation of MR-compatible bone screws as skull fiducials, showing the locations of the implanted screws. (C, D) Images obtained during the second surgery showing the exposed screws and measurement of their stereotaxic coordinates using a stereotaxic micromanipulator before the planned injection procedure. (E) Postoperative MRI scan showing the injection site within the targeted regions of the striatum.

During the second surgery, the animal was again positioned in the stereotaxic frame using conventional earbar-based alignment. Before craniotomy, the previously implanted screws were exposed, and the stereotaxic coordinates of each screw were measured using a stereotaxic micromanipulator (Figure 3C,D). These measured coordinates were compared with the fiducial coordinates predicted from the MRI scan using the registration methods described below. After fiducial measurement, the planned viral injection procedure was performed. A postoperative MRI scan was then acquired to assess the targeting outcome. Manganese was included in the injection solution to allow visualization of the injection sites on MRI (Figure 3E) Fredericks, Dash, Jaskot, Bennett, Lerchner, Dold, Ide, Cummins, Der Minassian, Turchi, Richmond and Eldridge (2020).

### 2.3. Registration and Error Analysis

#### 2.3.1. Offset Translation

We modeled conventional earbar-based zeroing as a pure translation. The estimated translation offset is:

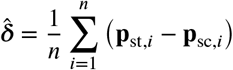

where *n* is the number of fiducials, **p**_sc,*i*_ is the coordinate of fiducial *i* in scan space, **p**_st,*i*_ is the corresponding coordinate in stereotaxic space, and 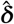 is the estimated translation offset. Scan coordinates were mapped to stereotaxic coordinates as:

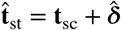

#### 2.3.2. Rigid Registration

We also evaluated rigid registration, which models the relationship between scan and stereotaxic coordinates as a rotation and translation without scaling or shear. In this approach, scan coordinates were mapped to stereotaxic coordinates as:

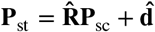

where **R** is a rotation matrix and **d** is a translation vector. The rotation matrix was constrained to preserve distances and orientation:

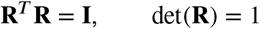

The rigid transformation was estimated from the corresponding fiducial coordinates by minimizing the squared error between predicted and measured stereotaxic coordinates:

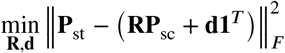

Using the estimated rigid transformation, target coordinates identified in scan space were converted to stereotaxic coordinates as:

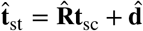

#### 2.3.3. Affine Registration

We also evaluated an affine registration method, which extends rigid registration by allowing translation, rotation, scaling, and shear. Unlike rigid registration, affine registration is not constrained to preserve distances or angles, but it can account for more general geometric differences between scan and stereotaxic coordinate systems. Affine registration was performed using homogeneous coordinates, so that translation could be included in a single transformation matrix. In matrix form, scan coordinates were mapped to stereotaxic coordinates as:

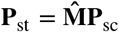

where **P**_sc_ and **P**_st_ contain the corresponding fiducial coordinates in scan and stereotaxic space, respectively, and 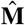 is the estimated affine transformation matrix. The affine transformation was estimated by:

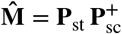

Using the estimated transformation matrix, target coordinates identified in scan space can be converted to stereotaxic coordinates:

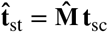

where **t**_sc_ denotes the target coordinates in scan space and 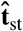 denotes the corresponding estimated coordinates in stereotaxic space.

#### 2.3.4. Registration Error Estimation

Registration error was quantified by comparing predicted stereotaxic coordinates with the corresponding measured stereotaxic coordinates. For each evaluated point, the residual error vector was defined as:

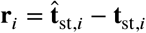

where 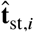 is the predicted stereotaxic coordinate and **t**_st,*i*_ is the measured stereotaxic coordinate of the *i*-th point. The mean absolute coordinate error for each evaluated point was calculated as:

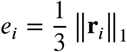

The average error across all evaluated points was then calculated as:

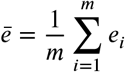

where *m* is the number of evaluated points. For independent target validation, *m* = 3, corresponding to three internal midline targets placed within the 3D-printed skull model. These targets were not used to estimate the registration transformations. Instead, their stereotaxic coordinates were predicted from the fiducial-based registration and compared with their measured stereotaxic coordinates.

In addition to registration error, linear correspondence between predicted and measured stereotaxic coordinates was assessed using axis-wise *R*^2^values. For each coordinate dimension, *R*^2^was calculated as the squared Pearson correlation coefficient between the predicted and measured coordinates. These values were used as a secondary measure of correspondence across coordinate axes, whereas mean absolute coordinate error was used as the primary measure of registration accuracy.

#### 2.3.5. Fiducial Resampling Analysis

To evaluate how the number of fiducials affected registration accuracy, the analysis was repeated using randomly selected subsets of *k* fiducials. For each subset size, fiducials were sampled without replacement, and the selected fiducials were used to estimate the registration parameters for each method: offset translation, rigid registration, and affine registration. The same fiducial subsets were used across registration methods and alignment conditions to allow direct comparison. For each value of *k*, the registration transform was estimated using only the selected fiducials. The resulting transform was then applied either to all fiducials or to the independent internal target points, depending on the analysis. Fiducial-based errors provided a measure of registration fit across the skull surface, whereas target-based errors provided an independent estimate of targeting accuracy because the internal targets were not used to estimate the transformations. To ensure bilateral coverage, fiducial subsets were selected from the left (*L*) and right (*R*) sides of the skull in a balanced manner for each *k*, such that:

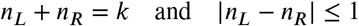

Registration errors were calculated as described above for each subset size and registration method.

#### 2.3.6. Asymptotic Registration Error

The asymptotic registration error was estimated by fitting the relationship between mean absolute coordinate error and fiducial number using a power-law decay model:

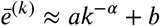

where *b* represents the asymptotic minimum error, and *a* and α describe the magnitude and rate of error reduction with increasing fiducial number. The model was fit using subset sizes of at least five fiducials because error estimates from smaller subsets were more variable and resulted in unstable power-law fits.

## 3. Results

We first evaluated the registration method using a 3D-printed skull model implanted with 28 MR-compatible screw fiducials. Three additional markers were placed along the midline inside the skull model to serve as internal targets, representing brain targets that were not used to estimate the registration transformations. After MRI scanning, the fiducials and internal targets were identified in scan space and measured directly in stereotaxic space. Using the matched fiducial coordinates, we compared three approaches under aligned and deliberately misaligned stereotaxic conditions (Figure 1). Offset translation was used as a reference model for conventional earbar-based zeroing, in which scan coordinates are shifted into stereotaxic space using a uniform coordinate offset but without correcting for rotation or other geometric differences. Rigid registration allowed both translation and rotation, while preserving distances and angles between points. Affine registration provided the most flexible transformation, allowing translation, rotation, scaling, and shear.

### 3.1. Registration Accuracy in the 3D-Printed Model

#### 3.1.1. Baseline Registration Accuracy

Under the aligned condition, all three registration methods established a correspondence between scan and stereotaxic coordinates. Linear regression between predicted and measured fiducial coordinates showed strong agreement across the three coordinate dimensions (*R*^2^> 0.98 for offset translation, *R*^2^> 0.99 for rigid registration, and *R*^2^> 0.99 for affine registration; Figure 2A). Across all fiducials, the mean absolute coordinate error was 0.47 mm for offset translation, 0.27 mm for rigid registration, and 0.27 mm for affine registration.

We also evaluated registration accuracy using the three internal midline targets, which were not used to estimate the transformations. For these independent targets, the mean absolute coordinate error was 0.28 mm for offset translation, 0.36 mm for rigid registration, and 0.65 mm for affine registration. The relative performance of the three methods as a function of fiducial number is described below (Figure 2B).

#### 3.1.2. Robustness to Misalignment

To further assess the robustness of each registration method, we introduced a misalignment condition by deliberately repositioning the 3D-printed model within the stereotaxic frame after MRI acquisition. This manipulation was intended to simulate positioning errors that may occur in practice, such as imperfect alignment between the earbars and ear canals or variability in surrounding soft tissue. Specifically, one earbar was removed from the ear canal of the 3D-printed model, and the fiducial and target positions were re-measured. The same registration analyses were then repeated under this condition.

The three registration methods differed in their sensitivity to stereotaxic misalignment. Under the misaligned condition, the linear correspondence between predicted and measured fiducial coordinates decreased for offset translation (*R*^2^> 0.88 across coordinate axes), whereas it remained strong for rigid registration (*R*^2^> 0.99) and affine registration (*R*^2^> 0.99). This indicates that transformations allowing rotational or more general geometric correction preserved the fiducial coordinate correspondence more effectively than offset translation alone.

Misalignment also increased the target error for offset translation, consistent with the limited ability of a pure translation to correct changes in head position. For the independent internal targets, the mean absolute coordinate error was 1.13 mm for offset translation, 0.42 mm for rigid registration, and 0.25 mm for affine registration. Thus, under the misaligned condition, rigid and affine registration maintained lower target errors than offset translation.

These results suggest that additional fiducials provide the greatest benefit for rigid and affine registration, while offset translation remains limited by its inability to correct rotational or other geometric differences. Given the 0.5-mm MRI voxel size, the target errors for rigid and affine registration were within approximately one voxel, indicating accuracy close to the spatial resolution of the imaging data.

#### 3.1.3. Effect of Fiducial Number on Registration Accuracy

Target registration accuracy generally improved as the number of fiducials used for parameter estimation increased. This pattern was observed across registration methods, although the magnitude of improvement differed between methods and alignment conditions. Offset translation showed limited improvement with increasing fiducial number, particularly under the misaligned condition, whereas rigid and affine registration benefited more from additional fiducials. The relationship between registration error and fiducial number approached an asymptotic minimum for each method. The power-law model fit the error curves well across registration methods and alignment conditions (*R*^2^> 0.98 for all cases; Figure 2C). This provided a quantitative basis for estimating the limiting registration error for each method and comparing performance across alignment conditions.

#### 3.1.4. Asymptotic Error Analysis

To summarize the relationship between fiducial number and target registration performance, we estimated the asymptotic error for each registration method by fitting a power-law decay model to the error curves. The estimated asymptotic error for offset translation increased from 0.26 mm in the aligned condition to 0.63 mm in the misaligned condition. For rigid registration, the corresponding values were 0.27 mm and 0.31 mm, indicating relatively stable performance across alignment conditions. For affine registration, the asymptotic target error was 0.64 mm in the aligned condition and 0.18 mm in the misaligned condition (Figure 2D).

We then compared the estimated asymptotic errors between aligned and misaligned conditions. Misalignment increased the asymptotic error for offset translation (*p* < 0.001), whereas rigid registration showed only a small change (*p* = 0.046). In contrast, affine registration showed lower asymptotic target error in the misaligned condition than in the aligned condition (*p* < 0.001). Across conditions, rigid registration provided the most stable target performance, with low asymptotic errors in both aligned and misaligned conditions. The higher target error observed for affine registration in the aligned condition may reflect partial overfitting to the skull-surface fiducials, because scaling and shear can improve the fiducial fit while reducing generalization to independent internal targets.

### 3.2. In Vivo Surgical Validation

As a proof of concept, we evaluated this approach in a rhesus monkey undergoing two surgical procedures. During the first procedure, ten MR-compatible screws were implanted in the skull and used as fiducial markers. As in the 3D-printed model, the most anterior point of each screw was identified in the MRI scan and used as the fiducial reference point (Figure 3A,B).

During the second procedure, the animal was positioned in a stereotaxic frame using conventional earbar-based alignment. After exposing the implanted screws, their stereotaxic coordinates were measured using micromanipulators and compared with coordinates predicted from the MRI scan (Figure 3C,D). Rigid registration also produced a stronger correspondence between predicted and measured stereotaxic coordinates (*R*^2^> 0.95 for offset translation and *R*^2^> 0.98 for rigid registration). The mean absolute coordinate errors were 0.55 mm for offset translation and 0.47 mm for rigid registration.

Postoperative MRI was used to assess the surgical targeting outcome (Figure 3E). Manganese was added to the viral solutions to allow visualization of the injection sites on MRI. The postoperative scan confirmed that the injections were localized near the intended targets, supporting the feasibility of using skull-fixed fiducials for MRI-guided stereotaxic targeting in vivo.

## 4. Discussion

In this study, we evaluated a skull-fixed fiducial-based framework for registering MRI coordinates to stereotaxic space in nonhuman primates. Using a 3D-printed skull model, we compared offset translation, rigid registration, and affine registration under aligned and deliberately misaligned stereotaxic conditions. Offset translation, used as a reference model for conventional earbar-based zeroing, was most sensitive to misalignment and showed limited benefit from increasing the number of fiducials. Rigid and affine registration made better use of multiple fiducials, but their performance differed depending on whether error was evaluated at skull-surface fiducials or independent internal targets. Affine registration provided a close fit to the fiducials, whereas rigid registration produced lower overall target errors, suggesting better generalization to internal locations. The in vivo validation further showed that implanted skull fiducials could be identified across surgical procedures and used to predict stereotaxic coordinates from MRI, supporting the feasibility of this approach for MRI-guided targeting in nonhuman primates.

The fiducial resampling analysis also showed that registration accuracy depended on the number of fiducials used to estimate the transformation. For rigid and affine registration, errors decreased as additional fiducials were included and approached an asymptotic minimum, suggesting that increasing fiducial number improves performance up to a practical limit. In contrast, offset translation showed less benefit from additional fiducials, consistent with the fact that a translation-only model cannot correct rotational or more complex positioning differences. These findings suggest that rigid registration may provide the most practical balance between correcting positioning differences and preserving anatomical geometry. A translation-only correction assumes that the difference between scan space and stereotaxic space is limited to a uniform shift, similar to conventional earbar-based zeroing, and therefore cannot account for rotations caused by small differences in head position. Rigid registration addresses this limitation by allowing rotation and translation while preserving distances and angles between the skull fiducials and internal targets. This constraint is important because the skull, implanted fiducials, and brain targets are expected to move together as a rigid structure. In contrast, affine registration can reduce fiducial error by allowing scaling and shear, but these additional degrees of freedom may partially overfit the skull-surface fiducials and reduce accuracy at independent internal locations. For practical targeting applications, rigid registration may therefore be a more conservative and reliable choice, especially when the goal is to predict small or deep structures that were not directly used to estimate the transformation.

The present method is closely related to the approach described by Bentley et al. (2018), who used skull-mounted titanium screw fiducials to improve stereotaxic targeting in nonhuman primates. Their study demonstrated that implanted cranial fiducials can provide a practical reference for MRI-guided procedures and showed the usefulness of this approach for targeting deep brain structures. Our study builds on this previous work in several ways. First, we systematically quantified registration accuracy under controlled conditions using a 3D-printed skull model. This allowed us to compare offset translation, rigid registration, and affine registration, examine the effect of stereotaxic misalignment, and measure how registration performance changed as the number of fiducials increased. Because the rigid transformation used here is similar in principle to the fiducial-based transformation used by Bentley et al., this comparison also allowed us to evaluate when additional model flexibility, such as scaling and shear in affine registration, is beneficial or potentially less reliable. These analyses allowed us to estimate the asymptotic registration error and evaluate the practical benefit of adding more fiducials. Another practical difference is the way the fiducials were visualized and localized in the MRI scan. In the previous approach, the titanium screws were identified mainly from the larger MRI signal void or artifact produced by the screw. In the present method, the MR-compatible screws were localized using a more specific fiducial point, such as the anterior point of the screw head. This may reduce uncertainty in fiducial localization and improve consistency across fiducials, particularly when the same implanted markers are used across imaging sessions and surgical procedures.

One practical limitation of this approach is that fiducials need to be implanted before the MRI scan, which requires an additional surgical step. However, this procedure is relatively simple and short, and the method uses standard MRI-compatible screws and conventional stereotaxic equipment. The approach also increases the duration of the stereotaxic surgery, because the fiducial positions must be exposed and measured accurately. Despite these practical costs, the in vivo validation showed that skull-fixed fiducials could be implanted, identified on MRI, and re-measured during a later procedure in a rhesus monkey. These findings support the feasibility of using implanted fiducials, particularly with rigid registration, for MRI-guided stereotaxic targeting across surgical sessions.

## 5. Conclusions

In summary, skull-fixed fiducials provide a practical reference for registering MRI coordinates to stereotaxic space in nonhuman primates. By using multiple fiducials and registration methods that account for both translation and rotation, this approach can reduce sensitivity to differences in head positioning between imaging and surgery. Although affine registration can provide a close fit to skull-surface fiducials, rigid registration may offer a more reliable balance between correction of positioning differences and generalization to internal targets. The results from the 3D-printed skull model and the in vivo validation suggest that this approach can improve the reliability of MRI-guided stereotaxic targeting, particularly for experiments that require accurate localization of small or deep brain structures.

## CRediT authorship contribution statement

**Peyton Harmon:** Methodology, Investigation, Formal analysis, Visualization, Writing – original draft, Writing – review & editing. **Reza Azadi:** Conceptualization, Methodology, Investigation, Formal analysis, Supervision, Writing – review & editing.

## Ethics approval and animal welfare

All experimental procedures adhered to the Guide for the Care and Use of Laboratory Animals and were reviewed and approved by the National Institute of Mental Health (NIMH) Animal Care and Use Committee.

## Acknowledgments

We thank the Section on Instrumentation at NIMH for their critical support, especially Katherine Cameron for her help with designing and 3D-printing the skull model. Anatomical MRI scanning was carried out in the Neurophysiology Imaging Facility Core (NIMH, NINDS, NEI).

This research was supported by the Intramural Research Program of the National Institutes of Health (NIH). The contributions of the NIH authors are considered Works of the United States Government. The findings and conclusions presented in this paper are those of the authors and do not necessarily reflect the views of the NIH or the U.S. Department of Health and Human Services.

## Appendix A. Supplementary material

Supplementary File 1: STL file of the 3D-printed skull model with fiducial locations used in the registration analysis.

## Data availability

Data will be made available on request.

